# Single chain models illustrate the 3D RNA folding shape during translation

**DOI:** 10.1101/2021.11.25.470027

**Authors:** Tianze Guo, Olivia L. Modi, Jillian Hirano, Horacio V. Guzman, Tatsuhisa Tsuboi

**Affiliations:** Department of Chemistry and Biochemistry, University of California San Diego, La Jolla, CA 92023-0358, USA; Department of Theoretical Physics, Jozef Stefan Institute, Jamova 39, 1000 Ljubljana, Slovenia; Institute of Biopharmaceutical and Health Engineering, Tsinghua Shenzhen International Graduate School, Shenzhen 518055, China

**Keywords:** RNA folding, RNA secondary structure, translation, modelling, bead-chain models

## Abstract

The three-dimensional conformation of RNA is important in the function and fate of the molecule. The common conformation of mRNA is formed based on the closed-loop structure and internal base pairings with the activity of the ribosome movements. However, recent reports suggest that the closed-loop structure might not be formed in many mRNAs. This implies that mRNA can be considered as a single polymer in the cell. We developed TRIP; *Three-dimensional RNA Illustration Program*, to model the three-dimensional RNA folding shape based on single-chain models. We identified the angle restriction of each bead component from previously reported single-molecule FISH experimental data. This simulation method was able to recapitulate the mRNA conformation change of the translation activity and three-dimensional positional interaction between organelle and its localized mRNAs. Within the analyzed cases base-pairing interactions only have minor effects on the three-dimensional mRNA conformation, and instead single-chain polymer characteristics have a more significant impact on the conformation. This method will be used to predict the aggregation mechanism of mRNA and long noncoding RNA in specific cellular conditions such as nucleolus and phase-separated granules.

## Introduction

RNAs are essential nucleic acids that convey genetic information, especially messenger RNAs (mRNA), which can be translated by ribosomes to produce proteins. RNA consists of single-stranded nucleic acid polymers that are connected to each other by forming phosphodiester bonds in the 5’ to 3’ direction. mRNA organization is important for many aspects of mRNA metabolism, particularly steps where different regions with (pre-) mRNAs are thought to communicate, such as splicing, translation regulation, or miRNA-mediated regulation (Fabian and Sonenberg, 2012; Imataka et al., 1998; Tarun and Sachs, 1996). However, despite the importance of mRNA organization, there is little known about how ribonucleoprotein complexes (mRNPs) are organized as three-dimensional assemblies. It has been observed that mRNAs exist in different levels of compaction depending on their translating state. These translating mRNAs are believed to exist in a closed-loop conformation where the 5’ and 3’ ends are brought together through the cap-binding elF4F complex and the poly(A) binding protein PABPC1 (Christensen et al., 1987; Imataka et al., 1998; Tarun and Sachs, 1996; Wells et al., 1998). Furthermore, since ribosomes will attach to mRNA during translation, this will add an unfolded and flat area to the strand, and the translation of mRNAs results in the separation of the 5’ and 3’ ends. This separation contentiously suggests that these RNAs are not translated in a stable closed-loop (Adivarahan et al., 2018).

In order to simulate the general pattern of RNA folding conformation, we adopted single-chain models and tested different angle restrictions. Such models were applied to break down the complicated structures of cytoskeletons and better understand aspects of the whole structure by considering smaller fragments (Wen and Janmey, 2011). Each element of the RNA strand can be categorized by the different links of a polymer-like arrangement of beads, and it simplifies the analysis of the whole function of the RNA. In addition, it draws a relationship between the effect of confinement on the end-to-end distances of single-chain polymers and the stiffness of each fragment (Doi, 1996). Even though this relationship is based on arbitrary numbers, it is able to characterize the generalized behavior and categorize the type of regime the single-chain falls into, which in turn allows an improved understanding of RNA conformation for the tackled cases and similar.

In this work, we applied the single-chain models for mRNA folding and developed the *T*hree-dimensional *R*NA *I*llustration *P*rogram (TRIP), a simplified model designed to rapidly interpret the end-to-end distances. The 5’ to 3’ distance of mRNA when treated with Puromycin, a known antibiotic that prematurely ends translation and disassembles polysomes, follows a Gaussian distribution with a skewness to the right (Adivarahan et al., 2018). Furthermore, ribosome occupancy determines the compaction of mRNAs; and the number of ribosomes positively correlated with the length of the mRNA (Adivarahan et al., 2018). When comparing our initial calculations with this single molecule Fluorescent in situ hybridization (smFISH) study, our findings gave an in-silico demonstration of this conclusion by simulating a chain of mRNA with and without ribosomes attached. Three-dimensional conformation of the mRNA is assembled by using a bent angle range of ±140°. Our single-chain models also recapitulated the distance between cytoplasmic mRNA and the mitochondria, which were previously analyzed in fixed cells by smFISH (Jourdren et al., 2010), and in live cells through an MS2-MCP method (Tsuboi et al., 2020). We also found that integration of the information of the local secondary structure, as it is the case of the tackled MS2-sequence, does not have much influence in the single chain model. While there is little known about mRNA’s three-dimensional organization in the cell, our simulation introduces a first approximation on how polymer-like models can rapidly estimate end-to-end distances of mRNAs in the cellular condition. In a nutshell, our very coarse model shows being useful for interpreting mRNA experiments devising end-to-end distances of the fragment.

## Results

### Modeling 3D RNA folding by TRIP; *T*hree-dimensional *R*NA *I*llustration *P*rogram

mRNAs are biopolymers consisting of thousands of nucleotides connected to each other from the 5’ to 3’ ends. The RNA backbone is rotameric (Murray et al., 2003). For each residue along the RNA backbone, there are six angles (α, β, γ, δ, ε, ζ) that display the rotation of the six bonds in each nucleotide. This suggests that RNA molecules can adopt complicated shapes because of the different combinations of angles that may occur. The single-chain bead model can be utilized to simplify the biopolymer by disregarding the atomistic details and considering each monomer unit as a structureless segment with a given length defined by monomer units and angles between monomers. By modeling three-dimensional RNA folding using TRIP; *T*hree-dimensional *R*NA *I*llustration *P*rogram, we analyzed the end-to-end distances by taking into consideration two biopolymer’s states one during non-translating condition and the other during translational elongation. These two states are similar to the conditions in which cells are treated with and without puromycin, translational inhibitor, prior to the formaldehyde fixation for the *in vivo* experimental measurements of end-to-end distance by the smFISH. In TRIP, we assumed single nucleotides as discrete monomers connected to each other (Figure 1A). The monomer length was set to be 0.59 nanometers based on an average distance between two nucleotides (Liphardt et al., 2001). The confinement of the angles is applied to the polar angle θ, the angle projection referenced to the y-z plane, and azimuthal angle φ, the angle projection referenced to the x-y plane. The set of (θ, φ) is randomly chosen from a defined angle range. We placed the first bead, the head of a single-chain, at the origin of a spherical coordinate and let it be the 5’ end, and the next bead was randomly picked from the outer surface of the sphere with a radius of 0.59 so that the first monomer, as a vector in the coordinate system, is generated with a length and direction. Based on the first vector’s direction, the next vector was created by setting the first bead as the origin and the second bead’s direction depending on the first vector’s direction within a confinement degree range (Figure 1B). This procedure was reiterated to create a strand of RNA.

**Figure 1.**
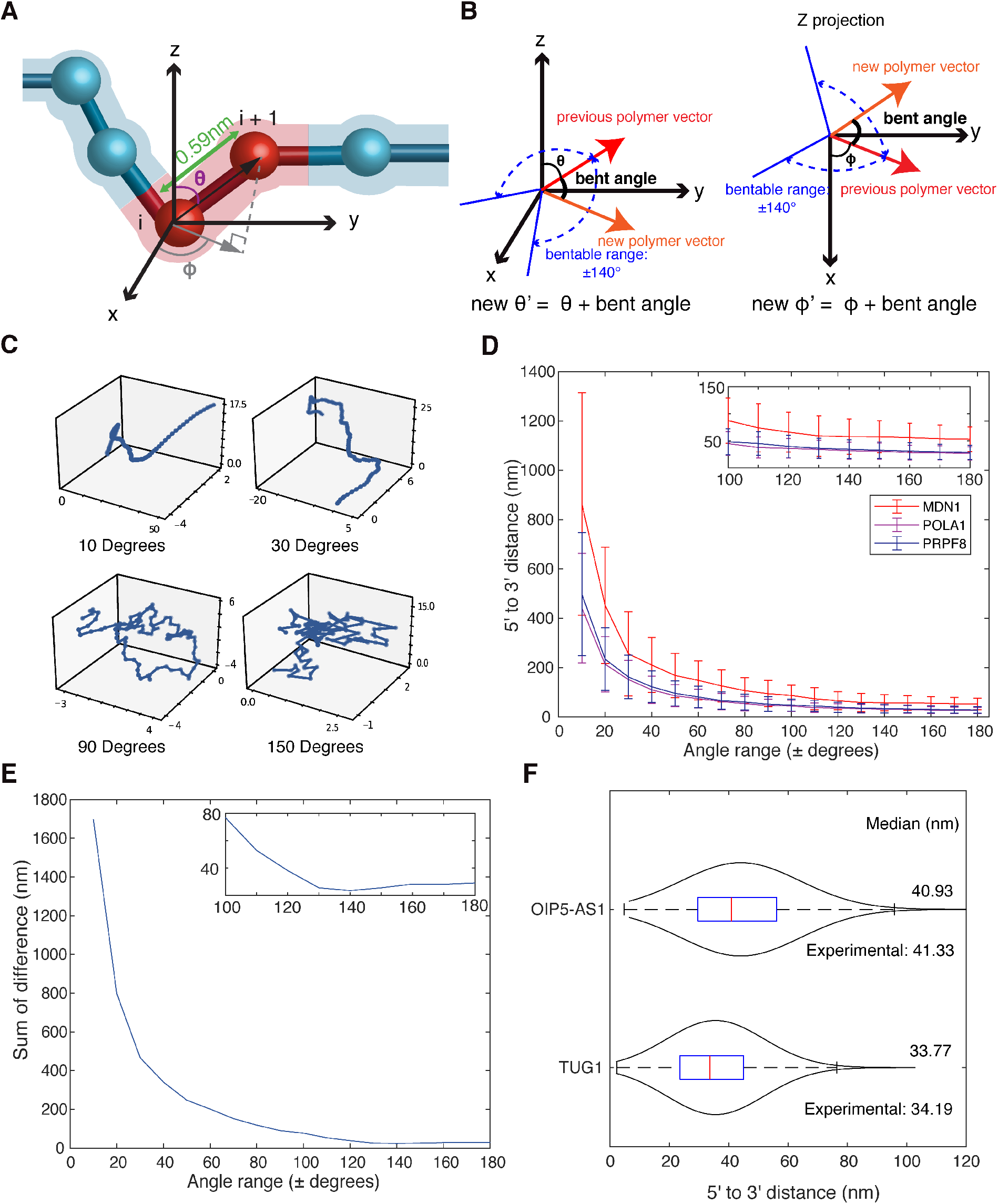
Simulating RNA 3D structure by RNA single chain models. (A) Diagram of TRIP; *T*hree-dimensional *R*NA *Illustration P*rogram, where the bead-chain in red is the area of focus. φ is the azimuthal angle, which is the horizontal angle between the single-chain and origin and measures the projection of the vector (indicated in black arrow) with reference to the +x axis. θ is the polar angle between the +z axis and the vector. The length between beads is 0.59 nm (Adivarahan et al., 2018). (B) The TRIP model is shown in two different projections, with three-dimensional Cartesian coordinates on the left, and Z projection on the right. The original vector is projected θ from the z axis and φ from the x axis, and the angle of the new single-chain vector is restricted by the 140 degree bendable range in both axes. (C) Visualization sample of 100 nucleotides RNA in the absence of ribosomes. The blue line indicates the length in nanometers the mRNA moves in each direction. The angle ranges are plus and minus 10, 30, 90, 150 degrees indicated respectively. (D) The relationship of 5’ to 3’ end-to-end distance simulated in two-dimensional with respect to the chosen angle range for *MDN1* (red), *POLA1* (magenta), and *PRPF8* (blue) mRNAs. Each dot indicates the median 5’ to 3’ distance on the given angle range, with error bars indicating one standard deviation. The frame on the upper right shows a zoom-in figure at angle range ±100-±180. (E) Plot of the sum of error, which is calculated as the sum of the distances of the TRIP outputs of three mRNA in (C) at a certain angle range to the previously reported smFISH data (Adivarahan et al., 2018). The frame on the upper right corner shows a zoom-in figure at angle range ±100-±180. (F) 5’ to 3’ end distance distribution of the non-translating RNAs *TUG1* and *OIP5-AS1* in two-dimensional. The blue box inside the violin plot shows the first quartile, median (red), and third quartile. The median from the simulation output is displayed on the right. The smFISH data (Adivarahan et al., 2018) is referenced below each plot.

While the monomer length was set, the angle range significantly affected the end-to-end distances. We used TRIP to visualize samples that were 100-nucleotides long and applied different angle ranges. It was observed especially at low angle ranges, such as 10 degrees, that the RNA strand will extend far; when angle ranges increase, the RNA strand tends to compact (Figure 1C, Movie 1). To identify the best angle range for TRIP, we compared the modified 2D TRIP results (Methods) to reported 2D 5’ to 3’ length for three different mRNAs: *MDN1, POLA1*, and *PRPF8* in non-translating conditions. Previous smFISH experiments tested the 5’ to 3’ distances in 3D and reported their result in 2D using image flattening techniques. The reported data were 35.95 nanometers (*MDN1*), 33.15 nanometers (*POLA1*), and 34.19 nanometers (*PRPF8*) respectively (Adivarahan et al., 2018). Because the measurement of length in smFISH is conducted by the distance between the centers of fluorescent signals from multiple probes, the nucleotides on the edge of the fluorescent signals may not be taken into account. To accurately input the experimental length for the simulation, we defined the net nucleotide number as the length between the centers of the FISH probes on the 5’ and 3’ ends. Different angle ranges were applied to the simulation for three mRNA references, with net nucleotides numbers 16350 (*MDN1*), 4060 (*POLA1*), and 4969 (*PRPF8*). As shown in the graph, there is a rapid decrease in the TRIP length first seen around ±10 to ±30 degrees, then at around ±100 degrees, the distances start to level off (Figure 1D). The results refer back to the observation that an increase in angle range leads to compaction. At around ±140 degrees the TRIP results showed high correspondence with the referenced values. To identify the best angle range, we analyzed the sum of the difference between the TRIP results and the previously reported smFISH data (Adivarahan et al., 2018) in each angle range (Figure 1E). By magnifying the larger angle range region, we confirmed that at ±140 degrees angle ranges, the sum of the difference is the least among all the angle ranges we tested (Figure 1E). We further confirmed if the ±140 degrees angle range could explain non-coding RNAs, which do not associate with ribosomes. Previous smFISH study showed that the non-coding RNAs’ end-to-end proximity was 41.33 nm for *OIP5-AS1* and 34.49 nm for *TUG1* (Adivarahan et al., 2018). The TRIP successfully recaptured these smFISH data as 40.93 nm for O*IP5-AS1* and 33.77 nm for *TUG1* respectively (Figure 1F).

### mRNA end-to-end distance positively correlates with numbers of ribosome attached

Ribosomes cover approximately 30 nucleotides of mRNA and unfolded and flattened mRNA during translation (Lareau et al., 2014). We tested if translation influences the mRNA conformation using TRIP. To analyze the relationship of translation activity and mRNA conformation, we used the 6636-nucleotide long SINAP mRNA, in which end-to-end proximity distance has been reported with a different number of ribosomal activities by smFISH (Figure 2B) (Adivarahan et al., 2018). The 5072 net nucleotides numbers were analyzed using fluorescent FISH probes at its 5’ and 3’ ends for this mRNA (Figure 2B). The open reading frame length of this mRNA was 4917 nucleotides long (Figure 2B). It has been shown that the maximum ribosome-bound number was 20 for 3129 nucleotides long open reading frame in SINAP mRNA previously (Wu et al., 2016). We estimated the maximum number of ribosomes for 4917 nucleotides open reading frame as 32 ribosomes by calculating the ratio. We conducted the *in silico* experiment for five groups, each with 0, 8, 16, 24, and 32 ribosomes attached, similar to what was shown in the FISH experiment. We observed a stepwise increase in length with evenly distributed intervals according to increasing numbers of ribosomes (Figure 2C). We also observed that the mRNA length distribution was a Gaussian distribution with a skewness toward the longer distance (Figure 2C). The relationship between distance and ribosome number suggests that more intense translation activity can increase the end-to-end distance, thus the molecule would be less compact since unfolded mRNA areas are greater.

**Figure 2.**
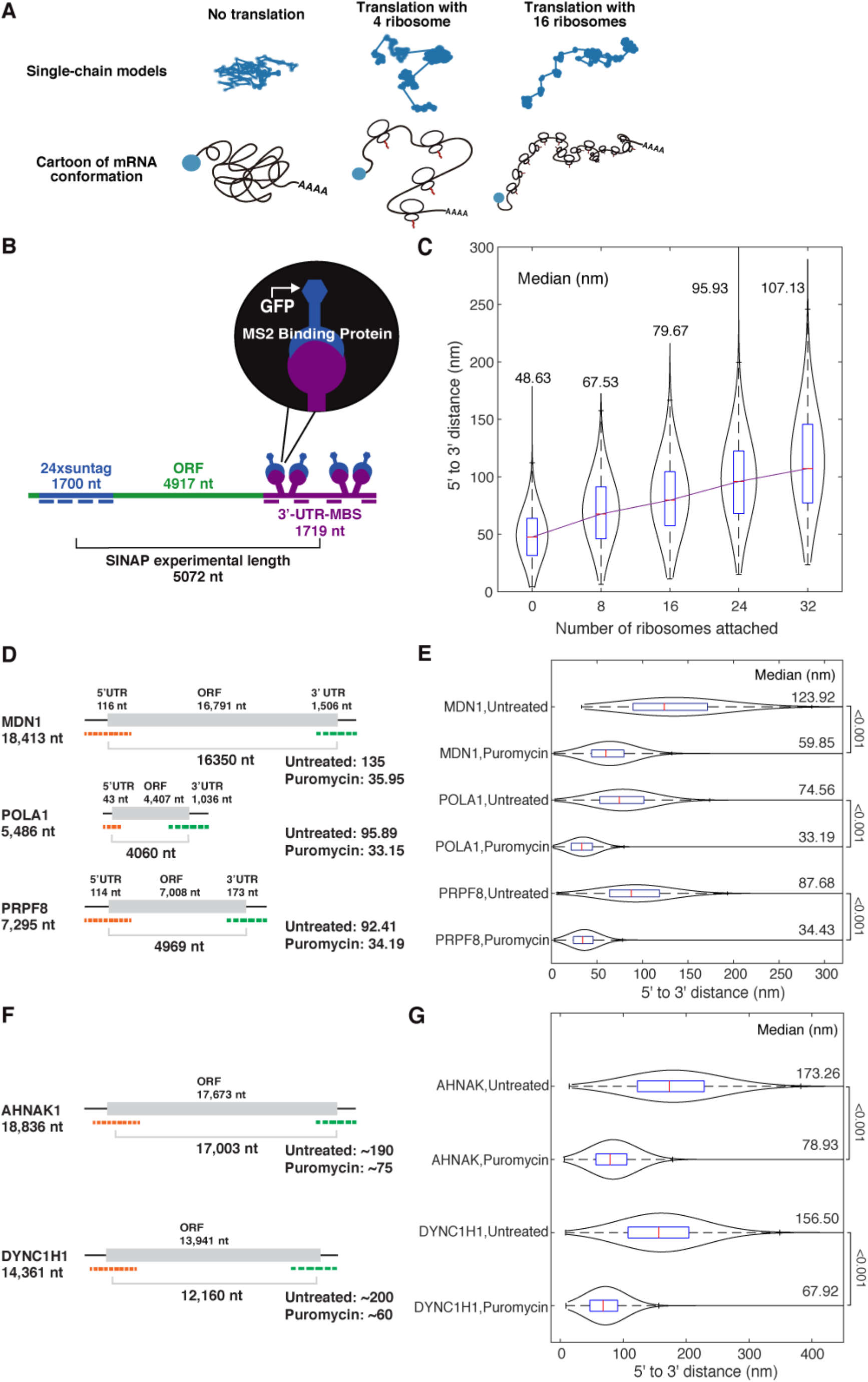
The ribosome occupation explains the separation of 5’ end and 3’ end of mRNA in different mRNAs. (A) Three-dimensional RNA Illustration Program models the three-dimensional RNA folding shape during no translation, mild translation, and intense translation (4 ribosome and 16 ribosome in the case of cartoons). (B) Flat model of the SINAPs mRNA structure and length. The blue dashed line represents 24xsuntag FISH probes. The purple dashed line represents 3’-UTR-MBS FISH probes. The experimental length is the absolute mRNA length between the centers of the above two probes (Adivarahan et al., 2018). (C) mRNA end-to-end distance positively correlates with numbers of ribosomes attached. Violin plots showing 5’ to 3’ distance distribution of the SINAP by TRIP simulation in the order of increasing translation activity in three-dimensional. The blue box inside the violin plot shows the first quartile, median (red), and third quartile. The median from the simulation output is displayed on the top. The purple line connects the data by their medians linearly. (D) Diagram of the mRNAs used in the previously reported experiments (Adivarahan et al., 2018). The bracket indicates the experimental length. The numbers indicated on the right exhibit the length in nanometers when cells are treated and untreated with puromycin in two-dimensional. (E) Violin plots showing 5’ to 3’ distance distribution of *MDN1, POLA1, PRPF8* mRNAs from the TRIP simulation for the cells treated and untreated with puromycin. The blue box inside the violin plot shows the first quartile, median (red), and third quartile. The median from the simulation output is displayed on the right. The 5’ to 3’ distance is simulated in two-dimensional. The number indicated by brackets outside the graph is the KS Test p-value between the two sets of data. (F) Diagram of the mRNAs used in the previously reported experiments (Khong and Parker, 2018). The bracket indicates the experimental length. The numbers indicated on the right exhibit the length in nanometers when cells are treated and untreated with puromycin in three-dimensional. (G) Violin plots showing 5’ to 3’ distance distribution of *AHNAK* and *DYNC1H1* mRNAs from the TRIP simulation for the cells treated and untreated with puromycin. The blue box inside the violin plot shows the first quartile, median (red), and third quartile. The median from the simulation output is displayed on the right. The 5’ to 3’ distance is simulated in three-dimensional. The number indicated by brackets outside the graph is the KS Test p-value between the two sets of data.

### The ribosome occupation explains the separation of 5’ end and 3’ end of mRNA in different mRNAs

We further tested if TRIP could recapitulate the impact of translation to end-to-end proximity distance using the three different mRNAs: *MDN1, POLA1*, and *PRPF8* mRNA referring to the previously reported smFISH data (Adivarahan et al., 2018). A significant difference of 5’ to 3’ proximity distance has been observed between the conditions of puromycin treated and untreated HEK293 cells (Figure 2D). These differences indicate that the translation contributes to the unfolding of mRNA conformations. We estimated the ribosome numbers as the average translational activity which is the length of the open reading frame divided by 200 nucleotides, where 200 is the average inter-ribosome distance on mRNA reported in human HEK293 and U2OS cells (Yan et al., 2016). Using this average number of ribosomes in all three mRNAs, we simulated the end-to-end proximity distance to compare with the previously reported smFISH results (Figure 2D) (Adivarahan et al., 2018). In all three cases, the TRIP predicted a significant difference between non-translating and translating conditions (Figure 2E). This result matched what was expected from in-vivo environments, in which the translation of these mRNAs resulted in the separation of the 5’ and 3’ ends (Adivarahan et al., 2018). The distribution of TRIP results follows a Gaussian distribution with a skewness to the larger distance, meaning a larger density was found at lower values. The compaction effect is found often in non-translating conditions, where the 5’ to 3’ end-to-end distance is significantly reduced from translating conditions. Interestingly, the end-to-end proximity for *POLA1* mRNA at translation condition (95.89 nm) was reported longer than *PRPF8* (92.41 nm) in the referenced data, despite *PRPF8* having a longer open reading frame (Adivarahan et al., 2018). However, the TRIP output predicted that *POLA1*-untreated is lower than *PRPF8*-untreated. This discrepancy might be due to *POLA1* possibly having higher translatability. We estimated that 40 ribosomes were required to reach the *POLA1* end-to-end proximity distance to the reported distance 95.13 nanometers by smFISH, which is 1.8 times greater translatability. In addition to comparing results shown in 2D space, we further investigated the ribosomal occupation on 3D data sets. Similar experiments using smFISH probes to detect end-to-end distance were performed to demonstrate the end to end distances of two sample mRNAs with long open reading frames under translating conditions and puromycin conditions: AHNAK (17673 nt ORF) and DYNC1N1 (13941 nt ORF) (Khong and Parker, 2018). The median is shown from their cumulative distribution plots as: AHNAK, translating conditions ∼190nm, and puromycin conditions ∼75nm; DYNC1H1, translating conditions ∼200nm, and puromycin conditions median ∼60nm. AHNAK and DYNC1H1 were simulated with net nucleotides numbers 17003 nts and 12160 nts according to the position of smFISH probes used in previous research (Khong and Parker, 2018). The TRIP output showed a similar trend when it was in 2D; it predicted a significant difference between non-translating and translating conditions (Figure 2F, G). Moreover, for AHNAK in translating conditions, the median is similar (173.26 nm) to the experimental data. For DYNC1H1 in translating conditions, the TRIP output shows a slight underestimation; the median (156.50 nm) is lower than the experimental data. This might be caused by the lower translatability of DYNC1H1. For both mRNAs in non-translating conditions, the TRIP outputs recapitulate the in vitro data: for AHNAK, the median gave 78.93nm; for DYNC1H1, the median gave 67.92nm (Figure 2G). In summary, the *in-silico* TRIP experiment was able to recapitulate that mRNA in a non-translation state follows a geometrically compact shape, whereas translation significantly separates the 5’ and 3’ ends.

### Local Secondary structure has minor effects in the studied cases

We tested how local secondary structures affect long mRNAs based on the lack of formation of circular structured mRNA. MS2 was used as a secondary structure in an effort to observe the way the mRNA folds. MS2 is a bacteriophage that consists of 3569-nucleotides of single-stranded RNA. It is encoded with four proteins: the maturation protein, the lysis protein, the replicase protein, and the coat protein (Fiers et al., 1976). The coat protein (MCP) is useful for the detection of RNA within living cells. For mRNA detection, MS2 binding sites (MBS) are inserted in the 3’UTR of an mRNA of interest, and the co-expression of MCP fused with fluorescent proteins renders single mRNAs visible using wide-field epifluorescence microscopy (Tutucci et al., 2018). Between the mRNA and MCP, there should be a secondary structure created by MS2. We predicted that by incorporating the information of the secondary structure, we would be able to refine the predicted RNA length. Each MS2 stem-loop structure was comprised of 17-nucleotides. This insinuates that the mRNA had an absolute experimental length of 1787-nucleotides and the structured mRNA length, which removed the 12 stem-loop sequences, was 1583-nucleotides (Figure 3A). It has been observed that there was a mean observed distance of 48 nanometers between the MCP-ORF in two-dimensional (Eliscovich et al., 2017). While the addition of this secondary structure could provide a refined predicted mRNA length, our analysis reveals that base-pairing interactions only have minor effects (2.3 nm for 204 nt) on the shape (Figure 3B), meaning that the effects of the MS2 secondary structure are less. This suggests that local secondary structure has minor effects on the tackled mRNA conformation.

**Figure 3.**
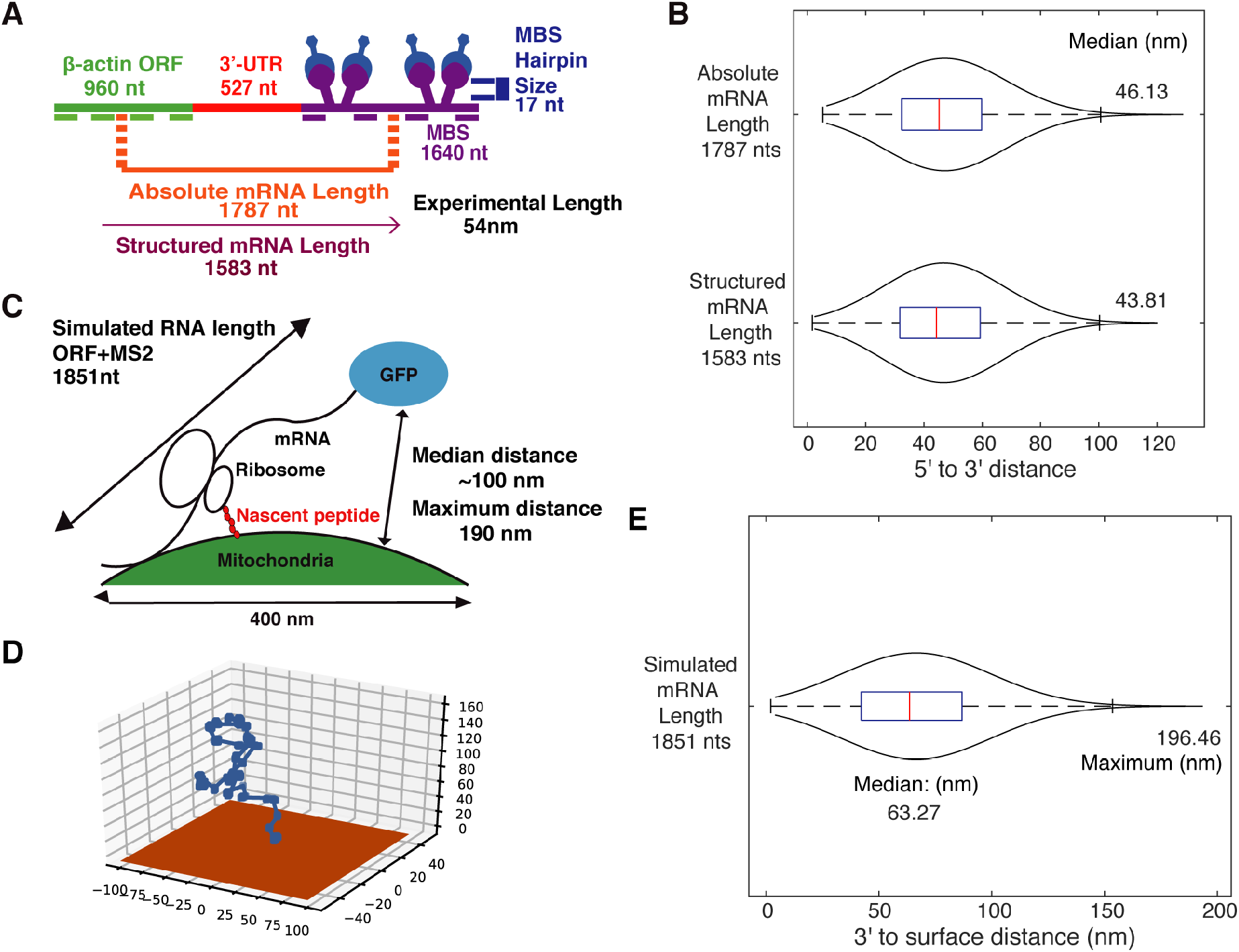
Determination of the conformation and length of a mitochondria localized mRNA. (A) Flat model of the experimental mRNA structure and length. The green dashed line represents ORF probes. The purple dashed line represents RNA FISH MBS probes. The structured mRNA length is the absolute mRNA length without the MS2 stem-loop sequence. Measurements for mRNA were taken in two-dimensional and obtained from Eliscovich et. al., 2017. (B) Violin plots showing 5’ to 3’ end distance distribution of the mRNA displayed in figure (A) with or without MS2 sequence in two-dimensional. The blue box inside the violin plot shows the first quartile, median(red), and third quartile. The median from the simulation output is displayed on the right. (C) Illustration of the distance between the mRNA and mitochondria in the previously reported experiments (Tsuboi et al., 2020). The indicated experimental distance was obtained from the association threshold between the mRNA and mitochondria. (D) Visualization sample of an mRNA located onto mitochondrial surface (orange surface), the beads indicate nucleotides and long edges stand for a ribosome has attached. (E) Violin plot showing 3’ end to the mitochondria surface distance distribution of the experimental mRNA displayed in (C). The blue box inside the violin plot shows the first quartile, median (red), and third quartile. The median and maximum distances from the simulation output are displayed on the graph.

### Determination of the shape and length of a mitochondria localized mRNA

Analyzing the proximity of nuclear-encoded mRNA to the mitochondrial outer membrane is challenged with long-term cycloheximide treatment by electron cryotomography (Gold et al., 2017). The distance between cytoplasmic mRNA and the mitochondria was also analyzed by visualizing single-molecule mRNA in fixed cells through smFISH (Jourdren et al., 2010), and in live cells using the MS2-MCP method (Tsuboi et al., 2020). We tested if the TRIP, which is a non-closed loop structure model, could recapture and explain the previous observations. To model such mRNA conformation, we defined the beads (i.e., nucleotides are not formed under the mitochondrial surface plane). To conduct the simulation, the x-y plane was set up as the “surface”, so that the result’s z-coordinate will never be less than zero. This represents a strand of mRNA growing on the x-y surface. A previous study with *TIM50* mRNA, a mitochondrially-localized mRNA (Figure 3C), has observed that the distance to the mitochondria follows a Gaussian distribution skewed to the larger distance, and the maximum distance is defined at 190 nanometers (Tsuboi et al., 2020). Using this information, we conducted the in-silico modeling by TRIP. 154 nucleotides were found as the average distance between ribosomes in budding yeast, *Saccharomyces cerevisiae* (Hurowitz and Brown, 2003; Zenklusen et al., 2008) and that *TIM50* mRNA’s open reading frame length is 1851 nucleotides. With this information, we estimated that 12 (=1851/154) ribosomes exist per mRNA. The visualization results of TRIP showed that a strand of mRNA tail-attached to the surface in orange color (*z* = 0) (Figure 3D). The TRIP results indicated the proximity distance between the mRNA 3’ and mitochondrial surface as a Gaussian distribution, with a skewness to the larger distance (Figure 3E). The median was at 63.27 nanometers and the maximum distance was 196.46 nanometers. The median value was far off from the experimental data (Tsuboi et al., 2020). It is possibly due to the inclusion of the non-associated mRNAs, which shows the distance of more than 190 nanometers, in the experimental data. The maximum distance was close to the previously reported experimental threshold value of 190 nanometers. This was also consistent with the previous FISH experiment showing the distance between mRNA localized to mitochondria and ribosomal RNA of mitochondrial matrix measured as 150nm to 200nm range (Jourdren et al., 2010). This suggests that the mitochondrial localized mRNA is also not forming a closed loop structure.

## Discussion

In this work, we presented that mRNA three-dimensional conformation can be approached as a simple single-chain bead model. We developed TRIP; Three-dimensional RNA Illustration Program and identified that 140 degrees is the restricted angle of each component of the single-chain to reproduce the previously reported experimental data set (Figure 1). We further tested the translational state by modeling the mRNA conformation to be unfolded and flat when the ribosome is associated (Figure 2). By analysing these sets of MS2 structure, we showed that the secondary structure, especially in the case of local and minor percentage of base paring in the mRNA fragments, would have a minimum effect on the mRNA end-to-end distance (Figure 3A, B). Lastly, we applied TRIP to predict mRNA conformation on the mitochondrial surface where it is experimentally difficult to address (Figure 3C-E). This methodology will be further applied to predict the aggregation mechanism for mRNA and noncoding RNA in specific cell organizations, such as the nucleolus and phase-separated granules.

The six rotatable torsion angles of the RNA backbone were analyzed through vector quantization (Hershkovitz et al., 2006), multiresolution approach (Hsiao et al., 2006), and quality-filtering techniques (Murray et al., 2003) and have met in consensus. The analysis of the six torsion angles together provides quantifications of the nucleic acids’ helical shape on an atomic level, and it also suggests that RNA can form various numbers of structures depending on the angles. However, at a coarser level this has to provide the torsion angle between the whole nucleotides. In our model we assumed each nucleotide position as beads, and instead of accounting for microscopic torsion angles between each atomic bond, we defined our set of bent angles (θ, φ) in a macroscopic view in which we assume and model each nucleotide broadly as observable clusters. The set of bent angles (θ, φ) explained that the relative position of each neighboring nucleotide is bent within 140 degrees positive or negative referring to the defined plane. We showed that an angle range of 140 degrees still gives a variety of RNA lengths based on our model. Compared to double helix structures like DNA, where each nucleotide is paired and forms a stable structure, RNA is relatively unstable and tends to compact or form secondary structures, thus, it may have smaller angles between each nucleotide. DNA on average has 10.5 bp/turn (Levitt, 1978), from geometrical analysis it suggests that each angle between nucleotides on average is approximately 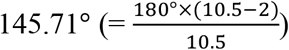. Our bent angle analysis, which was converted to an angle between nucleotides, ranges from 40°to 180°. Since our distribution was uniform in angle choosing, the statistical average angle between RNA was calculated to be 110°, which is less than that of double helix structures.

The evidence argues that ribosomes decompact mRNAs. An analysis of mammalian mRNP compaction by smFISH revealed that translating mRNPs are more extended compared to non-translating mRNAs (Adivarahan et al., 2018; Khong and Parker, 2018). The fact that mRNAs under stressed conditions show similar compaction compared to non-translating conditions may symbolize the removal of ribosomes (Khong and Parker, 2018). Translating mRNPs are compacted relative to its contour length (Adivarahan et al., 2018; Khong and Parker, 2018). This compaction is modelled by the formation of secondary structures of mRNA sequences between elongating ribosomes, which is supported by the spacing of ribosomes on mRNAs (Khong and Parker, 2020). Since the compaction of translating mRNPs is not affected by the inhibition of translation elongation, it is possible that stalled ribosomes on the open reading frame are sufficient to decompact mRNA (Khong and Parker, 2018, 2020). Due to spontaneous intramolecular RNA folding taking place rapidly, it can be assumed that mRNA sequences will collapse into RNA secondary structures, which are then unwound by each elongating ribosome (Khong and Parker, 2020). mRNA folding is also influenced by the binding of RNP-BP, including the ATP dependent binding of DEAD/Delhi box proteins (Khong and Parker, 2020). These box proteins, specifically DHH1, catalyse the ATP-dependent folding and remodelling of RNA duplexes. Immunoprecipitation assays reveal that DHH1 interacts with a variety of proteins, such as heat shock proteins, mRNA binding proteins, initiations and elongation translation factors, ribosomal proteins, and metabolic proteins (Marchat et al., 2015).

RNA’s secondary structure is described by the configuration of the base pairings that are formed by the biopolymer. These secondary structures can be divided into stems, which in naturally occurring RNA molecules are made up of five consecutive base pairs, and loops which connect or terminate the stems (Bundschuh and Hwa, 2002). While they locally form the same double helical structure as DNA molecules, RNA molecules are single-stranded, and therefore must fold back onto themselves to gain base pairings (Bundschuh and Hwa, 2002). The complexity of these secondary structures increase with length (Yoffe et al., 2008). We used the secondary structure formed by the MS2 bacteriophage, which is one of the simplest examples of a secondary structure. Our analysis showed MS2-structure only has minor effects on the shape (Figure 3B).

Intramolecular RNA-RNA base pairing interactions are important for small 5’-3’ distances. Previous research argued that the distance between the two ends of single-stranded RNA molecules is small, under the assumption that there are approximately equal proportions of A, C, G, and U (Yoffe et al., 2011). A large number of circle diagrams associated with the secondary structures of many RNA molecules was also examined for different lengths and sequences (Fang, 2011). It was revealed that the first and last monomers were always within a few monomers of each other, suggesting that it is essentially impossible to not have a minimum of one set of base pairs that will bring the ends of an RNA together, Furthermore, it was observed that certain RNA molecules, such as viral genomes (Poblete and Guzman, 2021; Poblete et al., 2021) and certain messenger RNAs, have been under selective pressure to maintain a small distance between the 5’ and 3’ ends of the molecule (Yoffe et al., 2011). However, this does not prove that biologically functional RNA molecules, such as viral genomes and certain messenger RNAs, have small 5’-3’ distance independent of sequence length (Clote et al., 2012). These findings were then experimentally confirmed by means of Förster resonance energy transfer (FRET). Theoretical analyses of a number of randomized and natural RNA sequences suggest that the 5’ and 3’ ends of long RNAs (1,000-10,000 nts) are always brought in the proximity of a few nanometres of each other regardless of RNA length and sequence because of the intrinsic propensity of RNA to form widespread intramolecular base pairing interactions (Lai et al., 2018). While secondary structures are thought to have a huge impact on mRNA three-dimensional conformation, this depends on the base-pairing percentage of the total mRNA fragment. In particular for the tackled biopolymers, our analysis would point towards base-pairing interactions only having minor effects on mRNA end-to-end distance.

## Supporting information

Movie 1 | Simulating RNA 3D structure by RNA single chain models

---

**Movie 1 | Simulating RNA 3D structure by RNA single chain models**

Visualization sample of 1-100 nucleotides RNA in the absence of ribosomes. The blue line indicates the length in nanometers the mRNA moves in each direction. The angle ranges are plus and minus 10, 30, 90, 150 degrees indicated respectively.

## Acknowledgments

We thank Dr. T. Wiryaman and members of the Zid laboratory for helpful discussions and feedback on the paper. We thank Dr. Y. Yang and Dr. B. Zid for critically reading the manuscript. This work was supported by startup funds from Tsinghua-SIGS (to T.T.). T.T. acknowledges support from a Japan Society for the Promotion of Science (JSPS) for a research abroad fellowship and postdoctoral fellowship (18J00995), and Uehara Memorial Foundation for research abroad fellowship. H.V.G. acknowledges financial support from the Slovenian Research Agency ARRS (Funding No. P1-0055).

## Author contributions

T.T. designed the study. T.G, O.M., and J.H. performed experiments T.G, O.M., and J.H. performed analysis. H.V.G. advised on the research. T.G, O.M., J.H., H.V.G., and T.T. wrote the manuscript. All authors discussed the results and commented on the manuscript.

## Data and materials availability

The code used for simulating the distance between 5’ and 3’ of mRNA and generating single-chain trajectory is available from https://github.com/paultianzeguo. Further information and requests for resources, scripts, and reagents should be directed to and will be fulfilled by the lead contact, T.T.

## Methods

### Modeling 3D RNA conformation

In this model, RNA molecules are considered single-chain consisting of multiple nucleotides and the atomistic details of each nucleotide were disregarded. Each nucleotide was considered as a structureless monomer of a given length r = 0.59 nanometers (Adivarahan et al., 2018). This model can be used to study a single-chain of beads with some prescribed restrictions between the monomers such as confinement of angles. We modeled the biopolymer by generating vectors that start from the origin, in three-dimensional Cartesian coordinates and shifting them to connect to beads with each other. The confinement of the angles is applied to the polar angle *θ*, the angle projection referenced to the y-z plane, and azimuthal angle *φ*, the angle projection referenced to the x-y plane. The set of (*θ, φ*) is randomly chosen from a defined bent angle range. The initial vector was randomly chosen as a given length r = 0.59 with a (*θ, φ*) set. The next angle was chosen based on both the previous (*θ, φ*) set and bent angle range.

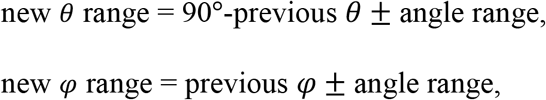

We repeated the process until the last nucleotide. In the process (*r, θ, φ*) was set and converted to (x, y, z), and added up to the previous vector in x, y, z coordinates. The conversion equation is:

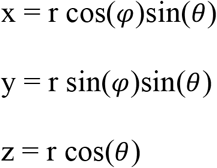

After repeating the addition of all nucleotides, the final output gave the vector (x1, y1, z1), which was converted to three-dimensional distance using 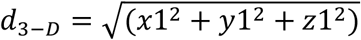. The experiment was repeated until it ran all the samples from a predefined sample number and graphed on a distribution plot. The sample number was chosen as 1000 for most of the analyses; for analyses on longer strands such as *MDN1*, 200 was used instead. For two-dimensional analyses, simple projection onto planes would have been biased; the spherical coordinates do not create a uniform length among the three possible projections onto the three planes. In the equation that converts the spherical coordinates to Cartesian coordinates, z was defined solely on the length r and the angle *θ*, however both x and y coordinates were dependent on the projection of r onto the z=0 plane, which is z= *r* sin*θ*, and the azimuth angle *φ*. Mathematically x and y components need to be multiplied by another trigonometry constant, which is less than one; in most of the cases x and y will be lower than its z counterpart. So, when we obtained the two-dimensional distance, we applied the approximation equation 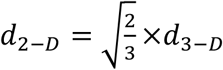, in which we approximate using an ideal condition when all the projections are the same, i.e., x1=y1=z1.

### Simulation of mRNA with ribosomes attached

In simulating mRNA with ribosomes attached, we assumed ribosomes had a 30-nucleotides long unfolded area. Based on the method used in simulating mRNA without ribosomes attached, we included a “add ribosomes” function after each strand of mRNA is formed. The resulting “mRNA” was a profile of coordinates that each nucleotide was in. We simulate the addition of ribosomes to let 30 continuous vectors in the same direction to be connected once, which can be simplified as elongating one vector to be 30 times its original length, and the rest adjust to this change in position. Before the simulation starts, the number of nucleotides should be adjusted to number of experimental nucelotide = number of nucleotides − 30∗ribosome number because the simulation of adding ribosomes counts as 30 individual nucleotides. After the resulting mRNA with the desired number of nucleotides is generated, the “add ribosome” function will run the number of times equal to the ribosome number set, in order to insert ribosomes. The function will randomly pick one point in the coordinate array and that point would be multiplied 30 times and the resulting value will be added to the rest of the nucleotides’ positions to adjust the following nucleotides’ positions. In order to avoid picking the same vector and multiplying it by 30 again, the function would include a checking process that determines if the place already had a ribosome attached to it. After all the ribosomes have been added, the function will read the last point’s coordinate (x1, y1, z1), and calculate its three-dimensional distance using 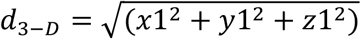 or two-dimensional distance as the same process as the mRNA simulation without ribosomes.

### mRNA on mitochondria surface

For simulating a mRNA strand on a surface, the x-y plane was chosen as the plane of reference. The simulation would start from the origin. We prevented the growing strand from going through the surface, i.e., *z* < 0. In order to accomplish this, we included an inspection process that checks if their z value is greater than zero after each new set of coordinates is formed.

### Visualization of the 3D mRNA model

All three-dimensional mRNA model visualization graphs were generated by the python package mplot3d from matplotlib. In this simulation, we collected the array of x, y, and z values as the loop function runs. It stores it in an array and uses it as input to the function mplot3d. All the other data-representing graphs and figures are generated in MATLAB from Mathworks.

