## Supplementary figures and images for "Single chain models illustrate the 3D RNA folding shape during translation"

### Movie 1 | Simulating RNA 3D structure by RNA single chain models

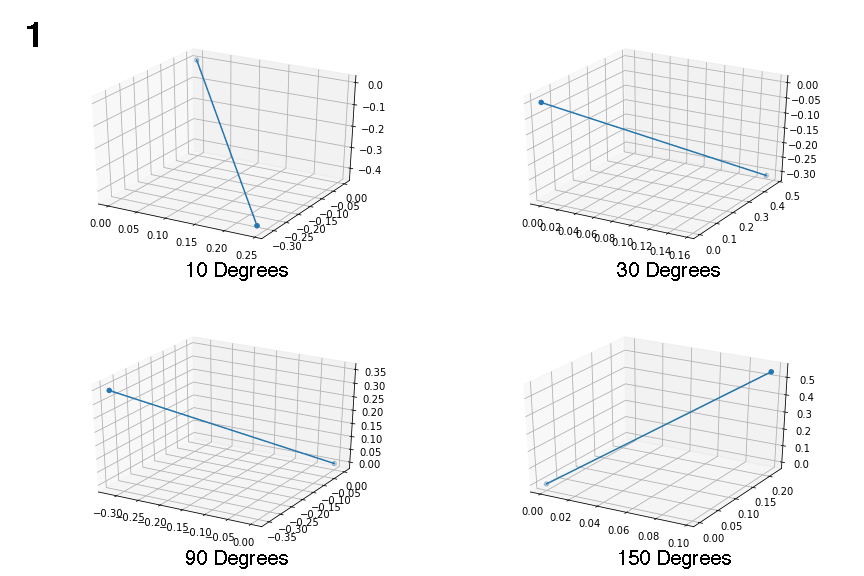
